# The Human T-cell Leukemia Virus capsid protein is a potential drug target

**DOI:** 10.1101/2024.09.09.612167

**Authors:** Ruijie Yu, Prabhjeet Phalora, Nan Li, Till Böcking, David Anthony Jacques

## Abstract

Human T-cell Leukemia Virus type 1 (HTLV-1) is an untreatable retrovirus that causes lethal malignancies and degenerative inflammatory conditions. Effective treatments have been delayed by substantial gaps in our knowledge of the fundamental virology, especially when compared to the closely related virus, HIV. A recently developed and highly effective anti-HIV strategy is to target the virus with drugs that interfere with capsid integrity and interactions with the host. Importantly, the first in class anti-capsid drug approved, lenacapavir, can provide long-acting pre-exposure prophylaxis. Such a property would provide a means to prevent the transmission of HTLV-1, but its capsid has not previously been considered as a drug target. Here we describe the first high-resolution crystal structures of the HTLV-1 capsid protein, define essential lattice interfaces, and identify a previously unknown ligand-binding pocket. We show that this pocket is essential for virus infectivity, providing a potential target for future anti-capsid drug development.

## Introduction

Human T-cell Leukemia Virus type 1 (HTLV-1) was discovered in 1979 and was the first retrovirus identified to cause disease in humans^1^. Approximately 10% of HTLV-1 carriers develop an aggressive and fatal malignancy called Adult T-cell Leukemia (ATL)^2^, and/or a progressive neurological inflammatory condition known as HTLV-1-associated myelopathy/tropical spastic paraparesis (HAM/TSP)^3^. The latter presents with debilitating clinical symptoms including muscle stiffness and weakness, bowel and bladder dysfunction and sexual dysfunction^4^. In addition to these incurable conditions, carriers may also present with other related inflammatory diseases, including pulmonary disease^5^, bronchiectasis^6^, uveitis^7^, and dermatitis^8^. With such a high-percentage of those infected developing cancer, HTLV-1 has been labelled the most potent oncovirus, and has even been argued to be the most carcinogenic substance on the planet^9^. Furthermore, there are currently no effective antiviral treatments available and prognosis for patients experiencing symptoms remains poor.

HTLV-1 is found across the world, with regions of endemicity on every inhabited continent. HTLV-1 has been classified into 7 subtypes (A to G), each resulting from zoonotic events that occurred over the last 60,000 years^10^. Subtype A, also known as the cosmopolitan strain, can be found around the world, and is especially prevalent in Japan, where it is estimated to have infected 1.1 million people^11^. Subtypes B, D, E, F, and G are largely restricted to Africa; while subtype C, also known as the Australo/Melanesian subtype, is found in Fiji/ Solomon Islands, Papua New Guinea and the indigenous peoples of central Australia. Indeed, these communities have the highest rates of HTLV-1 in the world, with recent studies suggesting as many as 40% of the adult population are positive for the virus^12^. HTLV-1c is the most divergent subtype, with estimates suggesting it separated from the rest 40,000-60,000 years ago and entered Australia about 9000 years ago^10,13^.

The other retroviruses known to cause disease in humans are HIV-1 and HIV-2, which are responsible for the AIDS pandemic. Over the last four decades, global efforts have seen the realisation of 7 drug classes for the successful treatment of HIV/AIDS, with the most recently developed class being the capsid inhibitors. Studies on the HIV capsid have shown that, rather than being an inert ‘shell’, the capsid orchestrates many of the early stages of viral infection. During the early post-entry stages of the viral life cycle, the capsid protects the viral genetic material from nucleases, such as TREX^14^, and innate sensors, such as cGAS^15^; it engages motor proteins to navigate the cytoplasm^16^; it functions as a semi-permeable reaction chamber to facilitate the process of reverse transcription^17^; it mediates nuclear entry^18–20^; and it partitions into membraneless compartments within the nucleus to direct the integration of viral DNA into specific regions of euchromatin^21^. Remarkably, the capsid is formed from a single protein, CA, which assembles into a fullerene cone comprising ∼250 CA hexamers, and exactly 12 CA pentamers^22^. The stability of the assembled capsid is finely tuned as it must ‘uncoat’ to release viral DNA at the appropriate time and location within the nucleus^23^. The multifunctionality and metastability of the capsid means that CA mutation often comes at a significant fitness cost to the virus. This ‘genetic fragility’^24^ and the fact that capsid carries pockets for host engagement has made the HIV capsid an attractive drug target in recent years.

Lenacapavir, the first in class anti-capsid drug, was approved for use by the European Union (EU)^25^ and the U.S. Food and Drug Administration (FDA)^26^ in 2022 for heavily-treatment experienced adults with multi-drug resistant HIV infection. Due to its slow-release kinetics from the injection site and long clearance times, lenacapavir need only be administered twice yearly. As such it is being considered for use in pre-exposure prophylaxis and is the first antiviral to ever show zero infections in a phase 3 HIV prevention trial^27^.

Lenacapavir was designed based on X-ray crystal structures of the HIV CA protein^28^. It binds primarily to a hydrophobic pocket on the surface of the HIV CA N-terminal domain (CA_NTD_) formed between helices 3, 4, and 5. Additional contacts with helices 8 and 9 on the C-terminal domain (CA_CTD_) of a neighbouring CA monomer ensure the drug preferentially binds the assembled capsid lattice (CA monomer *K*_d_ = 2500 ± 500 pM; CA hexamer *K*_d_ = 240 ± 90 pM). HIV uses this site to engage with multiple host proteins including, Sec24C^29^, CPSF6^30^, and the phenylalanine-glycine (FG) repeat domains of the nuclear pore complex^18,30^. By binding to this site, lenacapavir not only inhibits host cofactor engagement, it also promotes aberrant capsid assembly and disrupts mature capsid integrity^28,31^. These multiple mechanisms of action explain the high potency of lenacapavir (EC_50_ = 23 pM) and serve to highlight the therapeutic value of targeting the retroviral capsid.

The HTLV-1 capsid is almost identical between subtypes, despite over 50,000 years of divergence. This remarkable degree of conservation suggests that the HTLV-1 capsid is similarly genetically fragile, highly optimised to the human host, and may also represent a viable target for the development of long-acting prophylactics. As a structural protein with no enzymatic activity, it is necessary to understand the structure of the HTLV-1 CA protein, its lattice formations, and interactions in order to accelerate drug development in this space.

## Results

### Ultra-high resolution of HTLV-1 CA_NTD_

Across the known *Orthoretrovirinae*, CA is a two-domain protein with HIV being the best characterised. In this study, we have focused on HTLV-1 subtype C, isolate Aus-GM, isolated from a 67-year-old Western Desert Indigenous Australian male in 2013^13^ (GenBank JX891478). The HTLV-1c CA shares 97% sequence identity with subtype A and 32% with HIV-1 M-group CA (Extended Data Fig. 1). In order to determine the optimal boundary between the N-terminal domain (NTD) and the C-terminal domain (CTD), we predicted the full-length HTLV-1 CA structure using AlphaFold2. The prediction gave high-confidence folds for the individual domains, which were not spatially correlated with each other due to an apparently flexible linker (Extended Data Fig. 2a). Based on residue-level AlphaFold confidence scores, pLDDT (predicted local distance difference test), we determined the junction between the domains to occur between residues S127 and A128. A construct expressing CA 1-127, produced well in *E.coli* and was purified to homogeneity in the absence of an affinity tag by a sequence of ammonium sulfate precipitation, anion exchange, and size exclusion chromatography (Extended Data Fig. 2b). The protein crystallised in space group P1, diffracted to 0.87 Å resolution (Table S1), and was solved by molecular replacement using the AlphaFold2 prediction as the search model. The atomic-resolution electron density map unambiguously revealed an N-terminal β-hairpin (residues 1-13) followed by 6 α-helices and an additional 3_10_ helix located between helices 4 and 5 (Fig. 1a). Structure alignment with the HIV-1 CA_NTD_ showed three distinct differences.

**Fig. 1.**
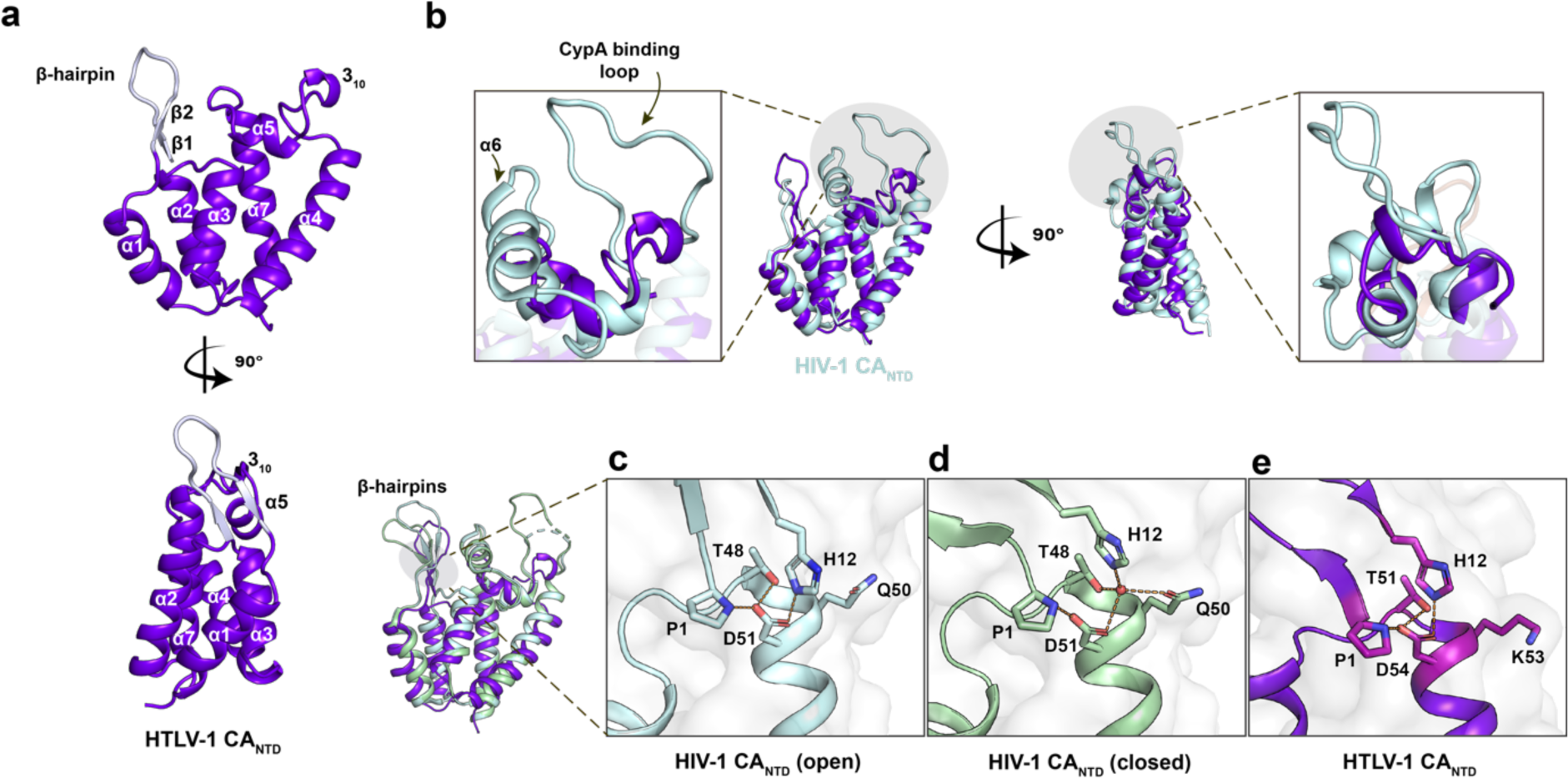
Structural comparison of HTLV-1 CA_NTD_ and HIV-1 CA_NTD_. **a**, The HTLV-1 CA_NTD1-127_ is labelled with secondary structure elements, including a β-hairpin (pale purple) formed by β-strands 1 and 2, α-helices 1-7, and an additional 3_10_ helix between α4 and α5, a 90° rotated view shown at the bottom. **b**, HTLV-1 CA_NTD_ is superposed onto HIV-1 CA_NTD_ (PDB 5HGL, in pale cyan). In HIV-1, the CypA binding loop and α6 are highlighted in grey shadow and magnified in the left box, a 90° rotated view shown on the right. **c-e**, HTLV-1 CA_NTD_ is superposed onto HIV-1 CA_NTD_ with open (PDB 5HGL) and closed (PDB 5HGN) states of β-hairpin, showing the residue interactions in the magnified views.

Firstly, HTLV-1 CA does not possess α-helix 6, which, in HIV, sits on the exterior surface of the capsid between the β-hairpin and the cyclophilin-binding loop (also known as the 4-5 loop, as it extends off the capsid surface between helices 4 and 5; Fig. 1b). In order to keep the nomenclature consistent between these structures, we have omitted helix 6 from our numbering (Extended Data Fig. 1).

Secondly, HTLV-1 CA is missing the extended 17-residue cyclophilin-binding loop, and instead has a shorter 11-residue sequence that adopts a more compact structure including a short 3_10_ helix. While a cursory inspection of the primary sequence alignment may suggest that residues 92-PLAGP-96 of HTLV-1 are similar to 90-PIAPG-94 of HIV-1, their structures do not align (Fig. 1b and Extended Data Fig. 1). Furthermore, the cyclophilin-binding loop of HIV CA has only been resolved when bound to either Cyclophilin A or Nup358^32^ as this loop is conformationally dynamic and contains the isomeric peptide bond (between G89-P90) that can adopt either *cis* or *trans* conformation. Conversely, the 4-5 loop of HTLV-1 CA is resolved in the absence of any bound cofactor and, while it contains two proline residues, all peptides are observed in the *trans* conformation.

Lastly, when HIV virions mature, the proteolytic release of CA from the Gag polyprotein results in an N-terminal proline residue which forms a salt bridge with a conserved aspartate sidechain (D51). The formation of this bond drives the folding of the N-terminal β-hairpin, which is a hallmark of CA maturity^33^. In some lentiviruses (including HIV-1 M-group, and SIV_cpz_), the β-hairpin has been shown to be able to switch between two states depending on crystallisation pH. In the open state (pH <7) H12 also forms a salt bridge with D51 (Fig. 1c), while in the closed state (pH >7) a water molecule is tetrahedrally coordinated between residues H12, T48, Q50, and D51^17^ (Fig. 1d). The consequence of these states is that the β-hairpin ‘opens’ or ‘closes’ like an iris above the six-fold axis of the CA hexamer. However, the residue at position 50 varies between lentiviruses and is frequently a tyrosine. Those viruses bearing Y50 (such as HIV-1 O-group, HIV-2 and SIV_mac_) are unable to bind the water molecule, cannot adopt the closed conformation, and consequently have a fixed ‘open’ conformation of the β-hairpin^15^.

HTLV-1 CA forms a structurally equivalent salt bridge between P1-D54 and a similar N-terminal β-hairpin (Fig. 1e). Residues P1 and H12 are well conserved, and T51, K53 and D54 are structurally equivalent to T48, Q50 and D51 of HIV-1 CA. In the HTLV structure, H12 contacts D54, consistent with the structure adopting the ‘open’ conformation (Fig. 1e). The presence of K53 suggests that HTLV-1 CA is not able to coordinate the water molecule necessary for adopting the closed conformation. We have solved the structure of the CA_NTD_ from both acidic and basic crystallants (see below and methods) and have observed no movement of the β-hairpin. We therefore conclude that the HTLV-1 CA β-hairpin is fixed in the ‘open’ conformation.

### Hexagonal crystal form of HTLV-1 CANTD reveals capsid lattice packing

In addition to our triclinic crystal form, we also obtained CA_NTD_ crystals in space group P622 that diffracted to a resolution of 2.05 Å (Table S1). This crystal form has one molecule in the asymmetric unit with hexagonal ‘sheets’ generated through crystallographic symmetry (Fig. 2 a,b). These sheets have features reminiscent of the HIV CA lattice packing observed in both crystallographic and cryoEM studies^34,35^, wherein the protein makes homotypic interactions to form a hexamer about the 6-fold axis (Fig. 2c). It is notable that the HIV CA_NTD_ is not capable of forming hexagonal lattices on its own, requiring the CTD to provide the necessary inter-hexamer interactions. In the case of the HTLV-1 CA_NTD_, inter-hexamer interactions occur at the crystallographic 3-fold axis, and are primarily mediated by residues from helix 4 (Fig. 2d).

**Fig. 2.**
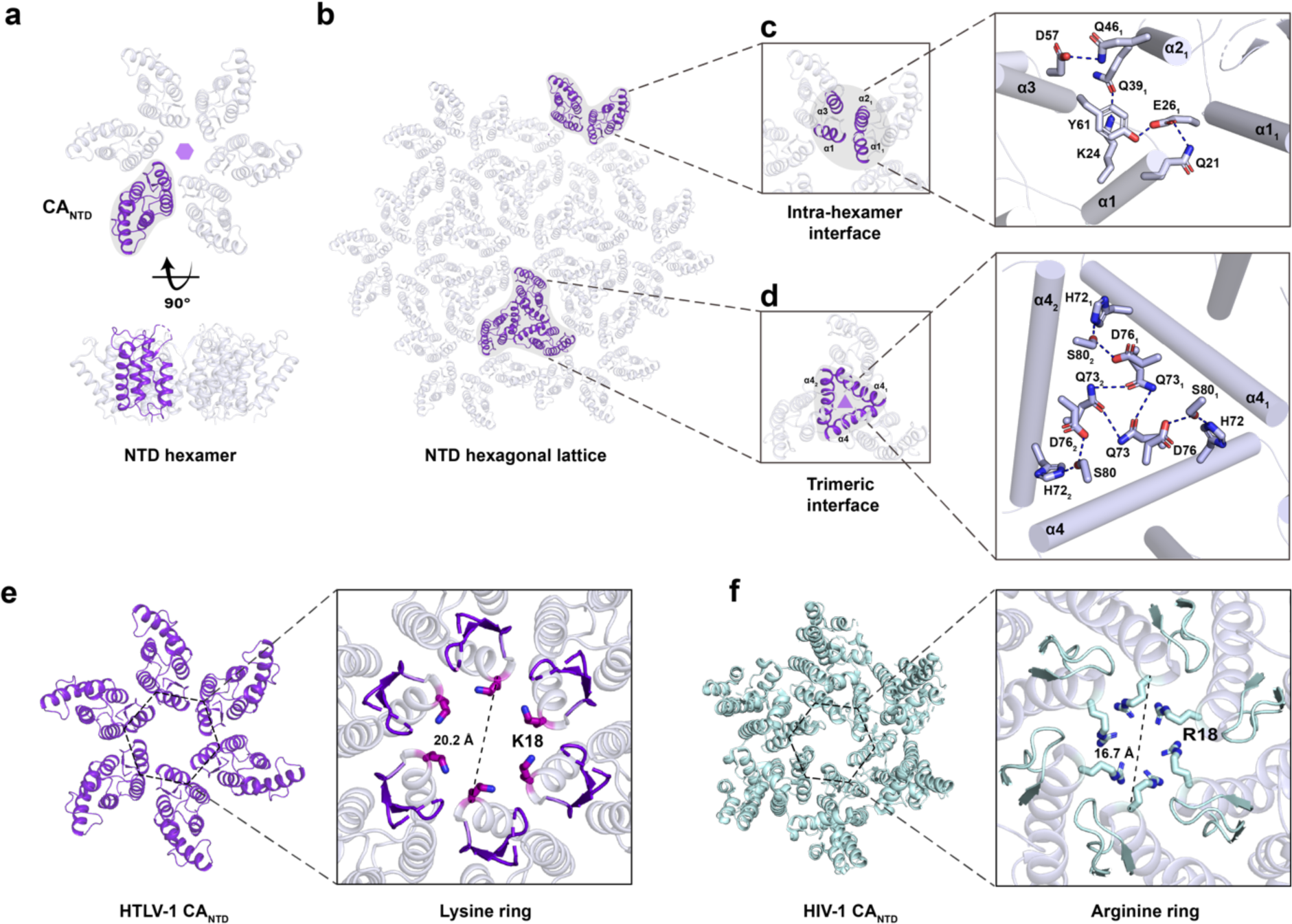
Features of HTLV-1 CA_NTD_ hexagonal lattice and interaction interfaces. **a**, Within the hexagonal crystal form, the HTLV-1 CA_NTD_ structure generates a hexamer that extends into a hexagonal lattice by crystallographic 6-fold symmetry (**b**). **c**, The intra-hexamer interface reveals the interaction between two NTD subunits, with relevant residues displayed as sticks. **d**, The trimeric interface among three inter-hexamers is formed by α-helices 4, with relevant residues shown as sticks. **e**, HTLV-1 CA_NTD_ hexamer contains six lysine residues at position 18, forming a positively charged pore at the centre of hexamer with a diameter of 20.2 Å. While HIV-1 CA_NTD_ hexamer (PDB 5HGL) has an arginine ring at position 18 with a diameter of 16.7 Å (**f**). HTLV-1 CA_NTD_ and HIV-1 CA_NTD_ are shown as the same scale, with diameter measured from C_α_ to C_α_.

Consistent with the mature HIV CA lattice, the HTLV-1 CA_NTD_ has a positively-charged ‘pore’ or ‘channel’ at the centre of the hexamer. This feature is formed by six symmetry-related lysine residues at position 18, which are structurally equivalent to the arginine 18 ring found at the centre of the HIV-1 CA hexamer (Fig. 2e,f). In HIV, the arginine pore has been shown to bind to polyanions such as inositol hexakisphosphate (IP6) which performs a lattice stabilisation function^36^, and deoxynucleoside triphosphates (dNTPs) that are the substrate for encapsidated reverse transcription^17^. It is likely that lysine 18 in HTLV-1 plays similar roles. We note that the pore is wider than HIV and attempts to co-crystallise with IP6 have not yielded compelling electron density at this site. As such we cannot rule out the possibility that HTLV-1 binds (an) alternative host factor(s) at this site.

### Full-length mature HTLV-1 CA crystallises as a hexagonal lattice

Previous studies on full-length HTLV-1 CA protein have reported difficulties producing soluble protein, which expressed poorly and degraded rapidly during purification. This was previously resolved by expressing a construct beginning at residue 16, effectively removing the N-terminal β-hairpin, and including an N-terminal His-tag^37^. We found that the full-length HTLV-1c CA protein expressed well in *E. coli*. Initial attempts to purify the protein by ammonium sulfate precipitation followed by cation exchange (at pH 5.0) were successful, but much of the protein precipitated when the pH was raised to 9 (note pI = 8.19). Surprisingly, inspection of the precipitate revealed that, rather than being amorphous aggregate, it comprised small clusters of thin hexagonal crystals. Screening buffer conditions by differential scanning fluorimetry showed a 6-degree increase in melting temperature when the pH was reduced from pH 9 (T_m_ = 54.6°C) to pH 6 (T_m_ = 61.3°C) (Extended Data Fig. 3a,b). The protein was therefore subsequently purified to homogeneity by size exclusion chromatography and stored in 20 mM HEPES (pH 7.0), 40 mM NaCl.

While readily crystallisable, only one condition yielded diffraction quality crystals after microseeding. The full-length HTLV-1 CA protein was solved to a resolution of 2.25 Å in space group F222 with three molecules in the asymmetric unit (Table S1). Applying a crystallographic 2-fold rotation generated the CA hexamer, packed as sheets, effectively identical to those observed in the hexagonal crystal form of the CA_NTD_ (Fig. 3a). Despite the high-resolution, electron density for the CTD was lower quality, suggesting that it remains flexible relative to the NTD in the crystal. Nevertheless, it was possible to trace the mainchain. To obtain higher resolution we expressed residues A128-L214 (CA_CTD_), which purified as a dimer (Extended Data Fig. 3c) and crystallised in space group P2_1_2_1_2_1_ with diffraction to 1.47 Å resolution (Table S1). The structure revealed a short 3_10_ helix followed by α-helices 8 to 11 (Fig. 3b, Extended Data Fig. 1). The asymmetric unit comprised 6 copies with two possible dimer configurations: one mediated by helix 9 (Extended Data Fig. 3d), and the other by an interdigitation of residues R188 and H190 in the 10-11 loop (Extended Data Fig. 3e). To improve the CA full-length structure, we re-solved it by molecular replacement searching for three independent copies of the refined CA_CTD_ structure as a search model. This unbiased approach returned a CTD dimer virtually identical to that mediated by α-helix 9 in the CA_CTD_ structure (RMSD = 1.16 Å, Figure 3c), suggesting that this is the biologically relevant dimer interface. This interface is centred on residue Y174. Curiously, while this residue is well-resolved in the electron density map from the CA_CTD_ structure, it makes an aromatic stacking interaction with its counterpart in the dimer partner that necessitates a breaking of the two-fold symmetry (Fig 3b). The observation that the CTD makes an unusual asymmetric homodimer may partly explain the difficulty in resolving this domain in the full-length structure.

**Fig. 3.**
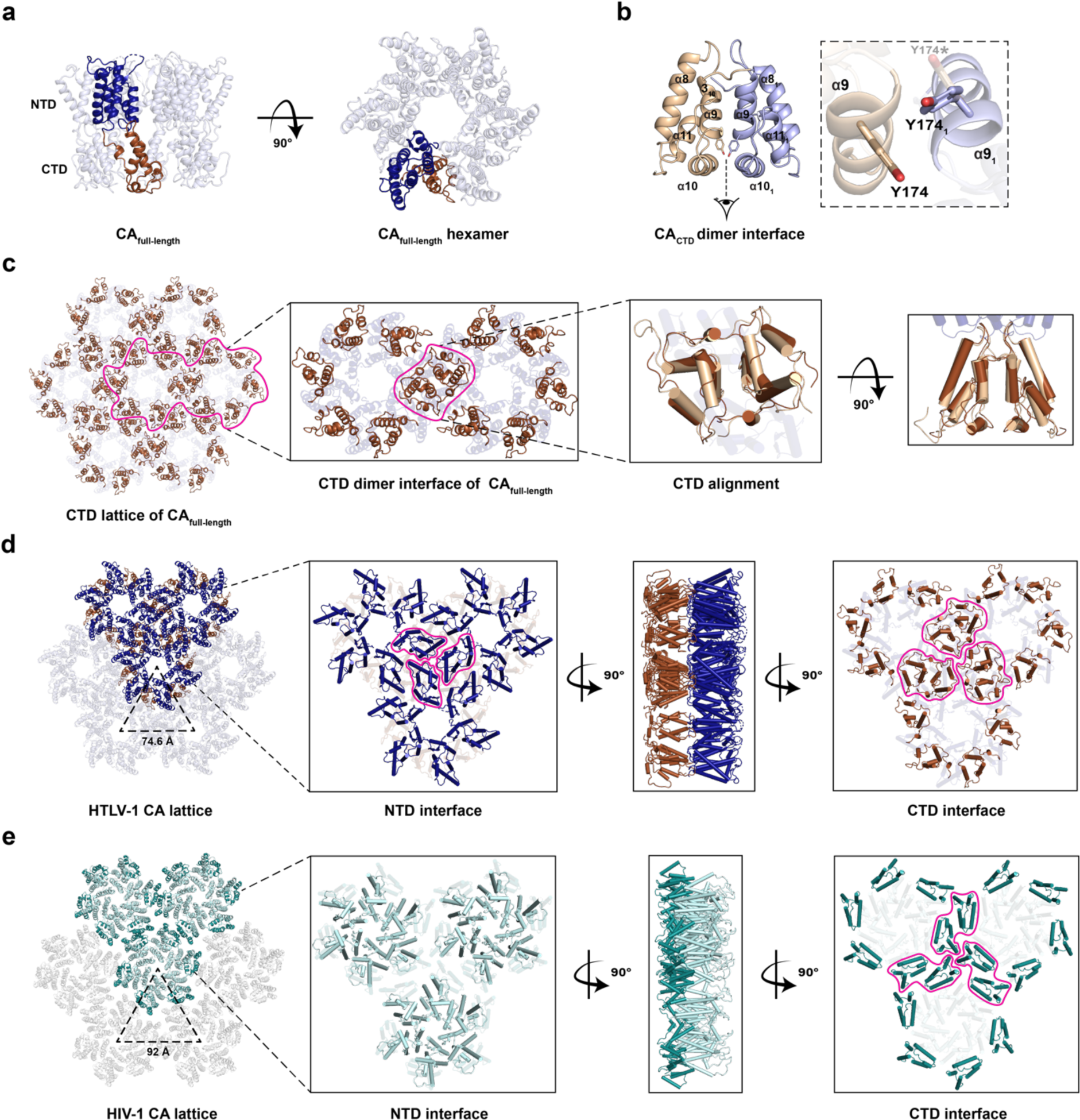
Comparing CA lattice differences between HTLV-1 and HIV-1. **a**, The crystal structure of HTLV-1 CA_full-length_ comprises domains of NTD (navy blue) and CTD (brown), shown in side and top views. **b**, The CA_CTD_ includes a 3_10_ helix followed by α-helices 8-11 and forms a homodimer centred on Y174. A bottom-up magnified view of Y174 shows that *π*-stacking with the dimer partner residue (Y174_1_, violet) breaks the two-fold symmetry. Y174* denotes the hypothetical symmetric conformation, and is shown for comparison with the actual asymmetric structure. **c**, The CTD of CA_full-length_ (brown) contributes to CA lattice packing by dimerising with a neighbouring inter-hexamer subunit, which aligns to the crystal structure of dimeric CA_CTD_ (wheat). **d**, The magnified interfaces of HTLV-1 and HIV-1 (**e**) CA hexagonal lattices are shown as a top (NTD face up), side and bottom (CTD face up) views with 90° rotation. The distance measured between two hexamer centres is 74.6 Å in HTLV-1 (**d**) and 92 Å in HIV-1 CA lattice (**e**).

The packing of the HTLV-1 CA into hexagonal ‘sheets’ is reminiscent of the lattice packing observed in other retroviral capsid crystal structures. In the case of HIV, the hexagonal lattice observed in crystals has been confirmed by cryo-electron tomography with subtomogram averaging^38^ to closely represent the hexagonal packing of CA within mature capsids. A key difference between the HIV (and SIV) and HTLV lattices is the nature of the inter-hexamer interactions. In the HTLV lattice, the NTD mediates both intra- and inter-hexamer interactions. The CTD is excluded from the NTD plane, but also contributes inter-hexamer interactions through CTD dimerization (Fig. 3d). This results in a closer-packed lattice for HTLV, with a distance between hexamer centres of 74.6 Å, nearly 20 Å shorter than that of the HIV lattice (92.0 Å). In the case of HIV, the CTD is in the same plane as the NTD and is solely responsible for the inter-hexamer interactions (Fig. 3e).

### Crystallant binding pocket may indicate a druggable site

During screening, a third crystal form of HTLV-1 CA_NTD_ was observed diffracting to 1.47 Å resolution in space group P2_1_2_1_2_1_ with two molecules in the asymmetric unit. In this case, the crystallant included 0.2M ammonium sulfate. In one of the HTLV-1 CA_NTD_ molecules, a sulfate ion is unambiguously resolved bound to an electropositive pocket on the NTD surface formed by α-helices 3, 4, 5 and 7 (Fig. 4a-c). Upon revisiting the lower-resolution P622 hexagonal crystal form, which also contained ammonium sulfate in the crystallant, a sulfate was observed bound at the same site. The sulfate ion is coordinated by residues H71, H72, R98, and W117 (Fig. 4a,c). The sidechains of Q56, H72 and R98 appear to act as gatekeepers that flip up and down to allow the sulphate entry into the pocket (Supplementary Video 1).

**Fig. 4.**
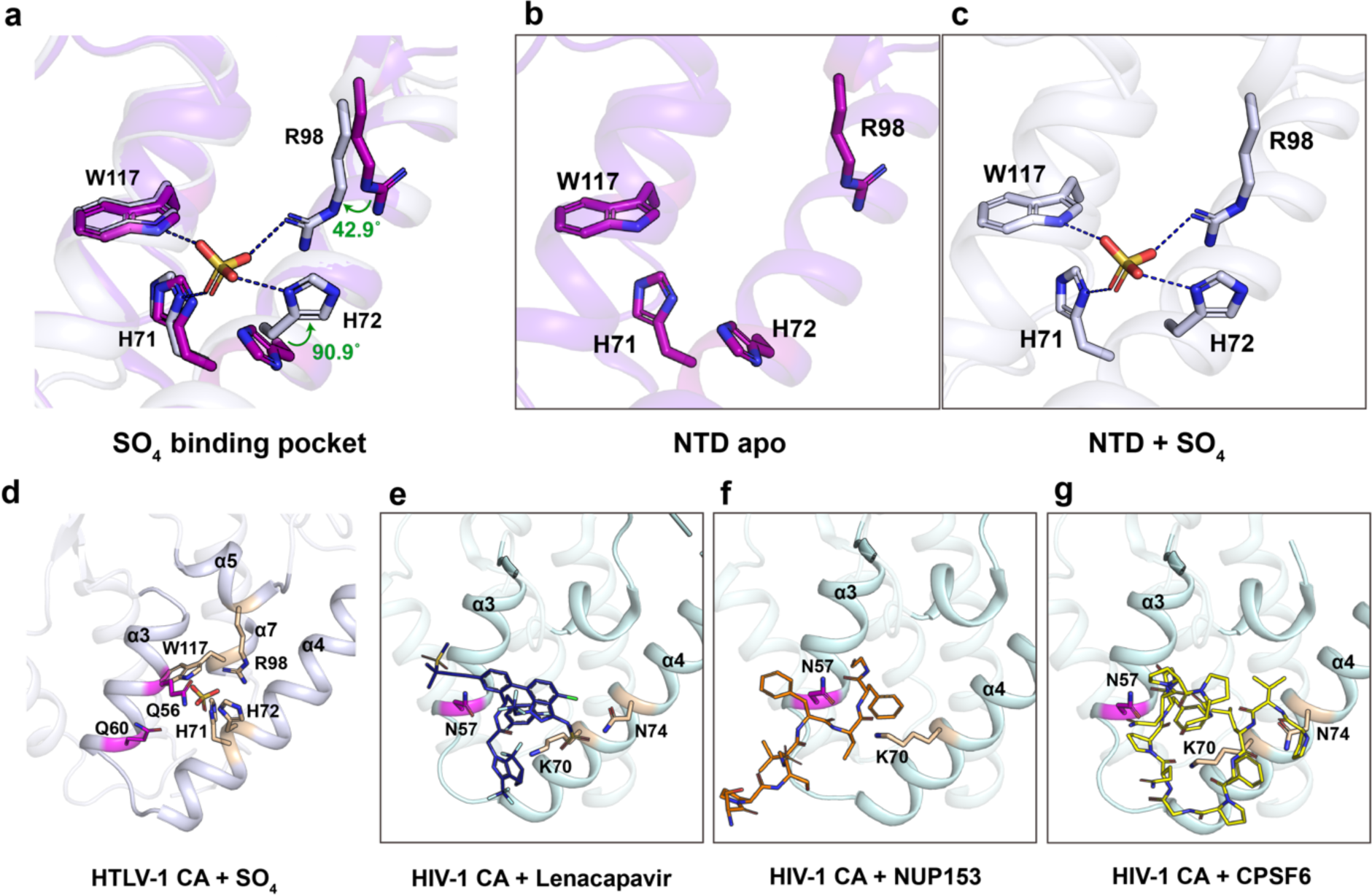
A SO_4_ binding pocket discovered in HTLV-1 CA_NTD_ orthorhombic crystal form. **a**, The structural superposition of apo CA_NTD_ (**b**) and SO_4_-bound CA_NTD_ (**c**) reveals that the SO_4_ binding site is enclosed by residues H71, H72, R98 and W117. **d**, The SO_4_ binding site is structurally equivalent to the FG-binding site in HIV-1 CA_NTD_. **e-g**, In HIV-1 CA, the FG binding site is occupied by lenacapavir (**e**, in dark blue), NUP153 (**f**, in orange) or CPSF6 (**g**, in yellow), respectively. Key residues interacting with ligands are highlighted in magenta and wheat.

In the HIV CA_NTD_, the equivalent pocket formed by helices 3, 4, 5 and 7 is hydrophobic and is responsible for recruiting host cofactors Sec24C, CPSF6, and FG-Nucleoporins^30,39^ via their phenylalanine-glycine motifs (Fig. 4e-g). Importantly, this site is the target of the first-in-class capsid inhibitor, lenacapavir^28^, which not only competes for cofactor binding, but also disrupts the capsid ultrastructure^31^. As this site has functional and therapeutic relevance to HIV, we sought to explore whether a biologically relevant compound may bind, and whether our system has the potential for use in ligand screening. The conditions producing the ultra-high resolution triclinic crystal form were free of sulfate resulting in an apo structure. Using a glutaraldehyde cross-linking strategy, we were able to soak the crystals in 2M lithium sulfate to obtain a sulfate-bound structure at a resolution of 1.71 Å (Extended Data Fig. 4a), demonstrating proof of concept that this triclinic crystal form is amenable to ligand binding while still maintain a respectable resolution. As sulfate is unlikely to be relevant to the HTLV infectious process, we performed a similar soak with sodium/potassium phosphate, obtaining a phosphate-bound structure to an almost identical resolution (1.73 Å) (Extended Data Fig. 4b). Interestingly, in crystallants at physiological pH, we observed partial occupancy of both sulfate and phosphate, while full occupancy was achieved under acidic conditions (pH 6 and below). This phenomenon likely reflects the protonation state of the binding pocket, which includes two histidine sidechains (H71 and H72) that typically gain positive charge under acidic conditions.

### Capsid surface residues mediate infectivity and/or particle production

Our structural data identified several surface residues that may make contacts essential for capsid lattice formation and/or engagement with external cofactors. Determining the importance of specific residues to the infectious process is complicated by HTLV-1’s requirement for cell-to-cell transmission, necessitating co-culture of producer and target cells. To overcome this limitation, we used a previously described replication-dependent infection system based on fluorescent protein reporter expression in HEK-293T cells^40^. Importantly, expression of the reporter is inhibited in producer cells and the fluorescent protein is only produced upon successful reverse transcription and integration in target cells and therefore reports only on cell-to-cell transmission. This single cycle, plasmid-based system also provides a facile means of screening CA mutants, but was originally developed for the study of HTLV-1 subtype A. For the purposes of this study, we replaced the CA gene with that from subtype C. Cell-to-cell transmission of HTLV-1 is notoriously inefficient, with typical cultures reported to yield 1-3% GFP-positive cells. To our surprise, the inclusion of the HTLV-1 subtype C CA protein consistently improved infectivity of the reporter virus by approximately 50% relative to the wild type subtype A (Fig. 5a).

**Fig. 5.**
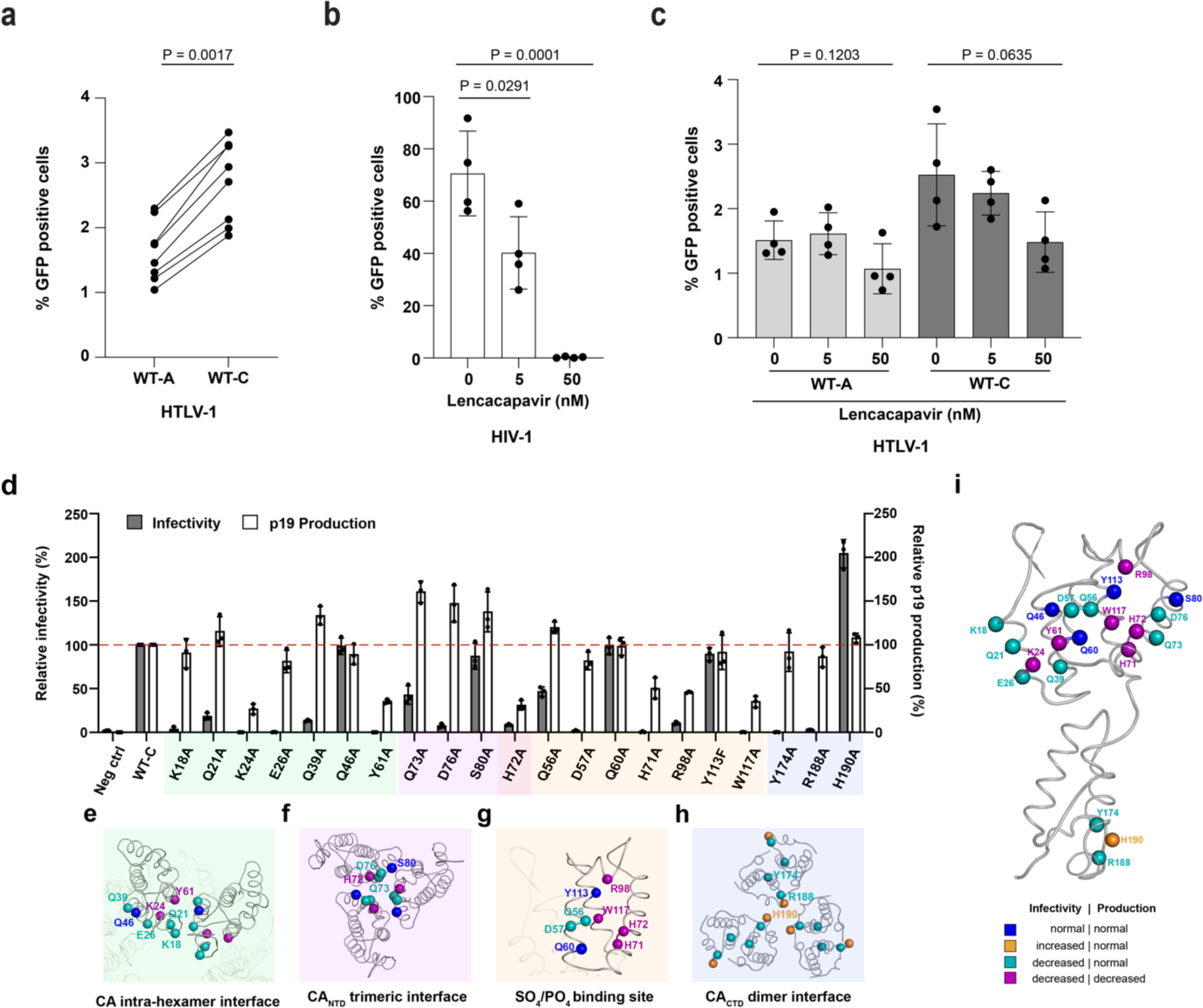
Infection and virus production phenotypes of HTLV-1c CA mutants. **a**, Graph comparing the infectivity of WT-A and WT-C packaging constructs as measured by GFP reporter expression in a replication dependent infection system. Results are displayed from 8 independent experiments. Statistical analysis was performed using an unpaired two-tailed t-test. **b** and **c**, Bar graph displaying infectivity of HIV-1 (**b**) and HTLV-1c (**c**) as measured by GFP reporter expression in a replication dependent infection system in the presence of increasing amounts of lenacapavir. Results are displayed as the average of 4 independent experiments with error bars representing the SD. Statistical analysis was performed using an unpaired two-tailed t-test. **d**, Bar graph displaying relative infectivity (grey bars) and p19 production (white bars) of 21 HTLV-1c CA mutants, compared to the WT construct, which is set at 100%. Shaded areas contain residues in the CA intra-hexamer interface (pale green, **e**), CA_NTD_ trimeric interface (pale purple, **f**), SO_4/_PO_4_ binding pocket (pale orange, **g**) and CA_CTD_ trimer of dimers interface (pale blue, **h**). Results are displayed as the average of 3 independent experiments with error bars representing the SD. **i**, HTLV-1c CA mutants, shown in **d**, mapped onto the structure of the monomeric CA full-length protein, with mutated residues coloured according to infection and virus production phenotypes.

Having established the infection model, we sought to test whether HTLV-1c is sensitive to lenacapavir in this system. While the equivalent HIV cell-to-cell infection was abolished at 50 nM lenacapavir, both HTLV-1 subtypes A and C were completely insensitive to drug treatment (Fig. 5b,c). Our results are consistent with recently published findings using replication-competent virus, which showed that lenacapavir treatment of either the HTLV-1a producer MT-2 cells or target Jurkat cells showed no discernible antiviral activity^41^.

To understand which surfaces on the HTLV-1 capsid might represent vulnerabilities targetable by future anti-capsid compounds, we developed a panel of 21 mutants on the HTLV-1c CA background. These residues were chosen based on their location within one or more of four putative interfaces: the intra-hexamer interface, the NTD trimeric interface, the CTD trimer of dimers interface, and the sulfate/phosphate binding pocket (Fig. 5e-h). Infectivity was measured by reporter gene (GFP) expression, while particle production was also assessed by p19 ELISA. All mutated residues within the intra-hexamer interface, with the exception of Q46A, substantially reduced viral infectivity. K24A and Y61A concomitantly reduced particle production by more than 50% (Fig. 5d,e), indicating that these two residues are likely indispensable for immature Gag lattice assembly prior to virus budding. Moreover, three mutations along α-helix 4 (H72A, Q73A and D76A), the main interacting interface in the CA_NTD_ trimer, significantly decreased infectivity (Fig. 5d,f), while S80A maintained wild type levels of infectivity and particle production. Mutants that would remove a direct interaction with the sulfate or phosphate groups (H71A, H72A, R98A and W117A) all effectively abolished infectivity with a concomitant loss of particle production (Fig. 5d,g). Importantly, it should be noted that, in the sulfate-bound state, H72 also spans the CA_NTD_ trimer interface (compare Fig. 2d and 4a). As such, it is conceivable that disruption of the ability of this site to accept a cofactor may have significant consequences for capsid lattice packing through disruption of the trimeric interface. In addition to those sidechains making direct ligand interactions, nearby residues Q56A and D57 in the sulfate/phosphate binding pocket were also found to be important for infectivity, albeit without impacting production, while mutants Q60A and Y113F were indistinguishable from wild type. Y113F was chosen in order to remove only the hydroxyl group of the sidechain, which binds to a water molecule bridging this residue to the sulfate/phosphate groups. Lastly, while we were confident in our assignment of the biologically-relevant CA_CTD_ dimer interface, we decided to test mutants at both this interface (Y174A), and the alternative interface suggested by the interdigitation observed in the CA_CTD_ crystal packing (R188A, and H190A – see Extended Data Fig. 3f). Unsurprisingly, Y174A reduced infectivity to zero, albeit without affecting particle production (Fig. 5d,h). R188A had an identical phenotype. Unexpectedly, H190A resulted in a 2-fold increase in infectivity compared to wild type, with no discernible effect on virus production (Figure 5 a). To the best our knowledge, this level of infectivity (∼7% GFP-positive cells; Extended Data Fig. 5a,b) is the most efficient HTLV-1 single cycle infection result to date.

Overall, we have identified four CA surface mutants phenotypically identical to wild type (Q46A, Q60A, S80A, and Y113F), ten mutants with decreased infectivity despite unchanged particle production (K18A, Q21A, E26A, Q39A, Q56A, D57A, Q73A, D76A, Y174A, R188A), five mutants with decreased particle production and infectivity (K24A, Y61A, H71A, H72A, R98A, W117A), and one mutant that increased infectivity (H190A) (Fig. 5i).

## Discussion

In this study we have focused on obtaining high-resolution crystal structures of the mature HTLV-1c CA protein. Despite previous studies reporting challenges with working with the HTLV-1a CA protein^37^, we found that the HTLV-1c CA and its constituent domains are easily expressed, highly stable proteins that routinely produced crystals surpassing the best resolution achieved for the more extensively studied CA from HIV^34,42,43^. We note that some of the residues that differ between subtypes A and C do participate in crystal contacts (such as T119 in the triclinic CA_NTD_ and T146 in the CA_CTD_ crystal forms), which may explain why subtype C is more amenable to crystallisation. Furthermore, in contrast to prior studies, our constructs carry the native N-terminal proline and β-hairpin (residues 1-13). As proline residues carry a positive charge only when found at the N-terminus, P1 can only form the salt bridge with D54 after CA is proteolytically released from Gag. As such, this salt-bridge and formation of the β-hairpin, both of which are well-resolved in our structures, are hallmarks of CA maturity. However, while it is likely that the hexagonal lattice packing observed in our full-length CA and CA_NTD_ crystals represent assemblies found within the HTLV-1 capsid, we have not formally demonstrated whether they represent the mature or immature packing. While the β-hairpin and the formation of the electropositive six-fold pore formed by residue 18, are both structural features uniquely found in the mature HIV lattice, a recent cryo-electron microscopy study performing subtomogram averaging on particles produced from the expression of immature HTLV-1a Gag^44^ produced a lattice similar to that found in our full-length CA crystal. In these particles, hexagonal CA packing is driven primarily by the NTD and the CTD is poorly resolved. It is possible that, in contrast to HIV, the HTLV-1 capsid undergoes a far less extreme structural rearrangement upon maturation. Future subtomogram averaging studies on the full-length mature capsid will provide further structural insight into the immature-to-mature transition, and possibly also reveal the structure of the CA pentamer, which has remained elusive to date.

As the two retroviruses that cause disease in humans it is useful to compare and contrast HTLV-1 and HIV. While both CA proteins share the same overall fold, there are significant differences, particularly at the capsid exterior. The fixed position of the HTLV-1c β-hairpin, the absence of helix 6, and the lack of a cyclophilin-binding loop (Fig. 1b) indicate that the cytoplasmic-facing surface of the CA_NTD_ is far less conformationally dynamic than that of HIV. If, like HIV, we assume that the HTLV-1 capsid acts as the host-pathogen interface during the early stages of infection, then it is clear that the two viruses engage with the host in different ways. One aspect where the two virus capsids are known to be functionally different is nuclear entry. Being a lentivirus, HIV is capable of penetrating the nuclear pore complex. It achieves this via specific interaction with FG dipeptide motifs found within the diffusion barrier in the centre of the nuclear pore^18,19^. Furthermore, it is likely disengaged from the nuclear pore complex by a competing interaction with a higher-affinity FG-motif in CPSF6^18,19^ which also likely directs the virus to the sites of integration. The hydrophobic FG-binding pocket is formed by helices 3, 4, and 5 on the CA_NTD_, and is also the site targeted by lenacapavir. By contrast, HTLV-1 is a deltaretrovirus and is incapable of entering the nucleus of resting cells, instead gaining access to host chromatin upon dissolution of the nuclear envelope during mitosis. As such, the HTLV-1 capsid does not have a hydrophobic FG-binding pocket and, consequently, it is insensitive to lenacapavir. However, our structures reveal that HTLV-1 does still have a pocket formed between helices 3, 4, and 5, albeit an electropositive one capable of binding to either phosphate or sulfate *in vitro*. While no capsid-binding cofactors have yet been identified for HTLV-1, it is conceivable that the pocket formed by helices 3, 4 and 5 is a more general ‘host recognition’ feature that has diversified over the course of retrovirus evolution. In the HTLV-1 case, sulfate is unlikely to be the endogenous cofactor, and inorganic phosphate is too ubiquitous to provide any spatiotemporal regulation to the capsid but could potentially function as a ‘pocket factor’^45^. A cryptic phosphopeptide, on the other hand, could conceivably function as a capsid-binding cofactor at this site. In HIV, N57 makes two hydrogen bonds with the main chain peptide of the phenylalanine of the FG-motif (see Fig. 4f,g). In HTLV, the structurally equivalent residue is Q60, which has similar hydrogen-bonding capabilities as the asparagine found in HIV. However, Q60A is phenotypically unchanged from wild type in our single round infection model. Alternatively, Q56 may function in a similar cofactor-recognition role.

Unlike FG-cofactor or drug binding in HIV, phosphate/sulfate binding in HTLV-1c results in significant structural changes, especially at residues H72 and R98 (Fig. 4 and Supplementary video 1). Mutation of the ligand-binding residues abolishes infectivity and reduces particle production (Fig. 5). One possible explanation would be that a bound cofactor is required for appropriate immature lattice formation prior to budding. It is particularly striking that H72 is so conformationally responsive to sulfate/phosphate binding, while also participating in the trimer interface, and also being essential for infectivity and particle production. As the equivalent site has been exploited for the development of arguably the most potent and longest-acting anti-HIV drug, our structural and phenotype data suggest that similar attention should be paid to this site in HTLV-1 with the view to the development of capsid-targeting pre-exposure prophylactics.

Across our panel, we identified ten surface mutants that reduce infectivity without affecting particle production. These mutants warrant further investigation as this phenotype could result from aberrant capsids and/or loss of an interaction or function important to infection. K18, for example, is the residue responsible for forming the electropositive pore at the six-fold axis, a feature found in all orthoretroviruses genera (with the exception of the gammaretroviruses). This residue likely performs a functional role (e.g. dNTP import) as we see no evidence for IP6 binding in the HTLV-1 case. Likewise, the discovery of a mutant that improves infectivity, H190A, is particularly interesting and also deserves to be the focus of future studies as this is an unusual phenotype to observe during alanine mutagenesis. Given its location in the CA_CTD_, it is not immediately apparent why H190A would improve infectivity. One possible explanation is that H190A improves the fidelity of mature capsid assembly. Given that the residue was identified due to its participation in a homotypic interaction (albeit a crystal contact), it is possible that inappropriate CTD:CTD interactions mediated by H190 occasionally occur during capsid lattice formation. Alternatively, H190A may enable HTLV-1 to escape from a cryptic restriction factor. In either case, the question remains as to why it is advantageous for the virus to maintain a histidine at this position given that its removal appears to enhance transmission. Nevertheless, the discovery that both HTLV-1c and H190A can improve the single round infection model raises the possibility that there may be additional ways to improve infectivity in this system (e.g. cofactor expression, restriction factor depletion, alternative mutants). If a single round infection model could be developed that could achieve higher rates of infection, this would enable high-throughput methods for cofactor identification (e.g. CRISPr screens).

To date there are no effective antiviral therapies to treat or prevent HTLV-1 infection. While HIV has been the great success story of antiviral development, it took nearly 40 years to realise that the capsid represents one of the best drug targets. The work we present here supports the notion that the same is likely true for HTLV-1: the capsid is vulnerable and has at least one pocket that could be exploited by a small molecule drug. The robustness of the recombinant CA protein and the ease with which high resolution crystal structures can be achieved supports the feasibility of developing a screening platform amenable to structure-based drug design. Furthermore, understanding the functions and interactions of the capsid during infection will undoubtedly shed light on how to best interfere with the spread of HTLV-1.

---

**Extended Data Fig. 1.**
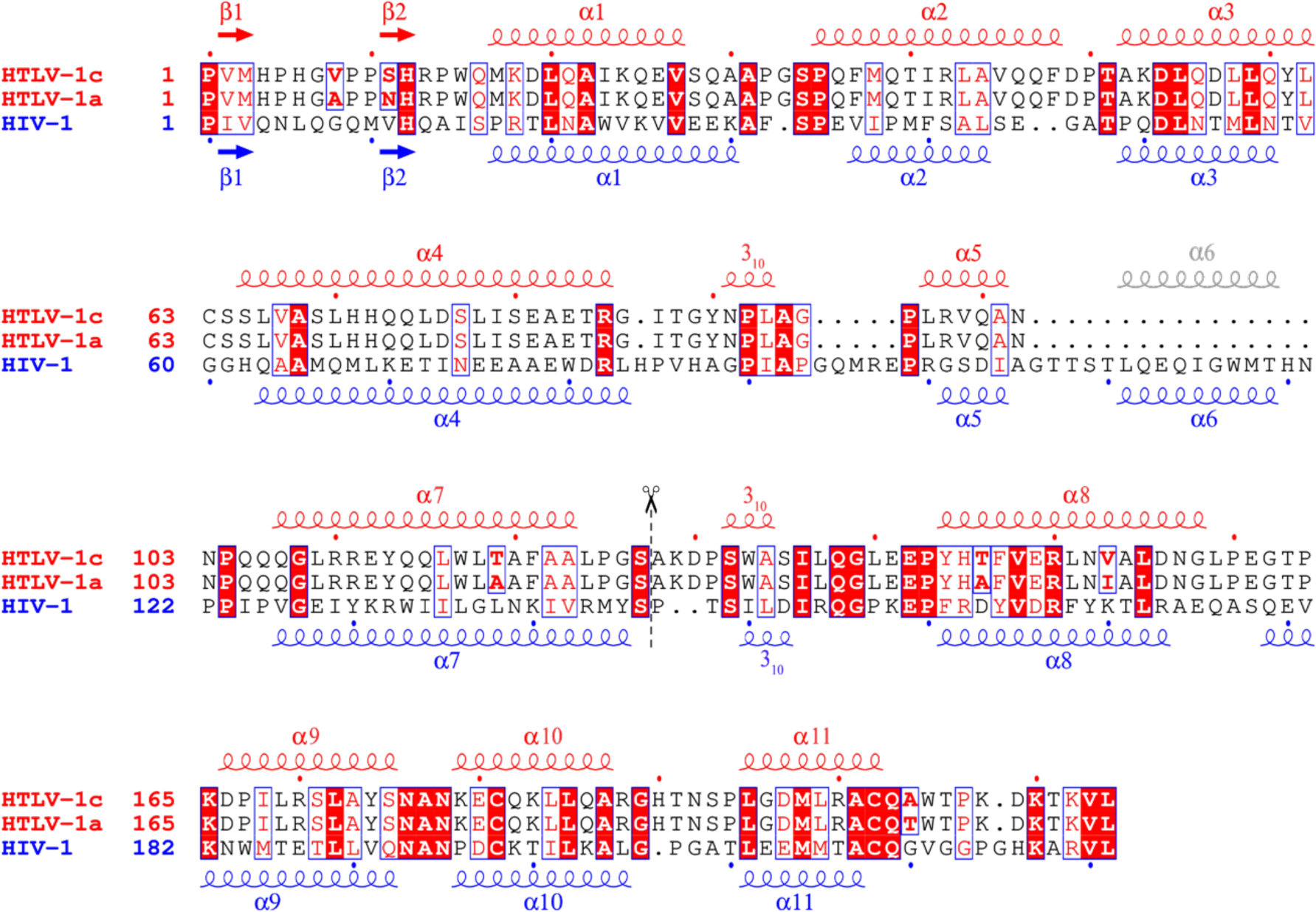
Structure-based sequence alignment of HTLV-1c, HTLV-1a and HIV-1 capsid proteins. The alignment is shown secondary structural elements of HTLV-1c/a CA (in red) and HIV-1 CA (in blue). Strictly identical residues are highlighted in red background, and the highly conserved residues are shown in red and boxed in blue. The alignment is generated using ESPript 3^46^.

**Extended Data Fig. 2.**
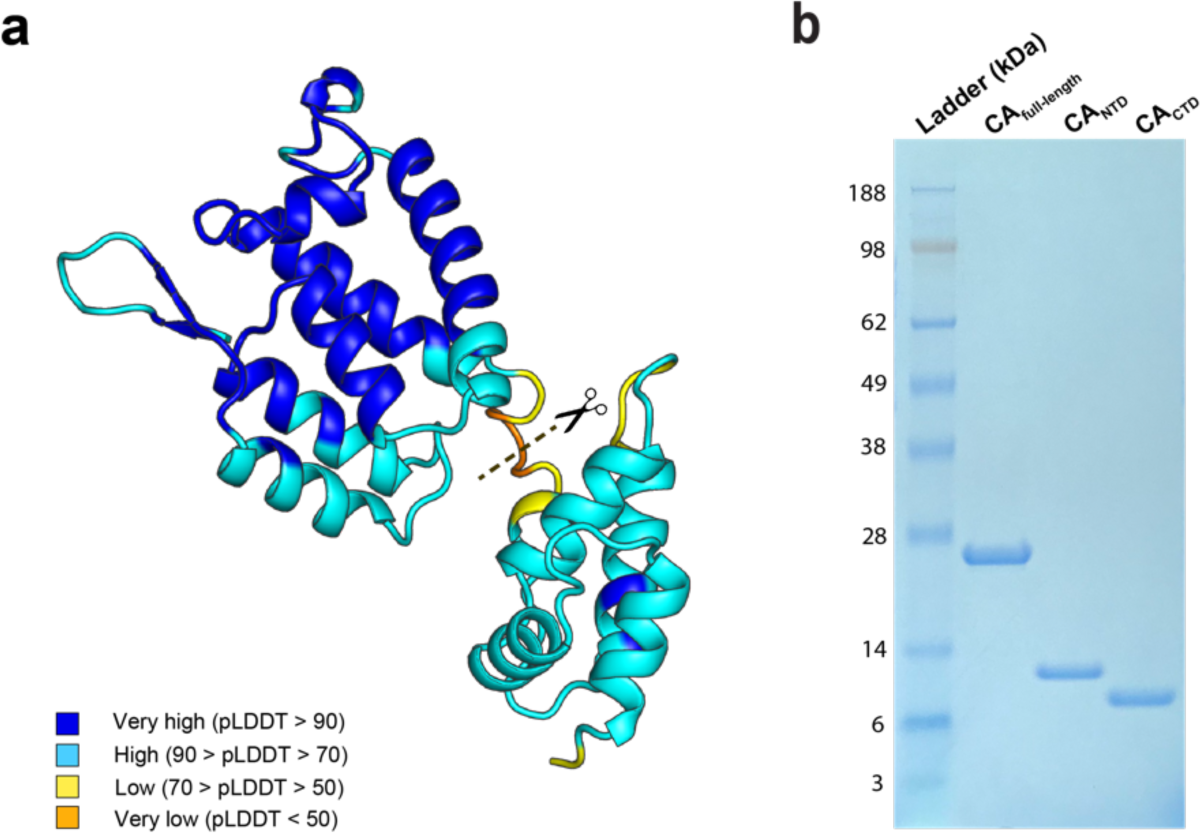
HTLV-1 CA model by AlphaFold2 prediction and the SDS-PAGE of CA proteins. **a**, The AlphaFold2 predicted model of HTLV-1 CA is coloured according to confidence levels, as measured by the predicted local distance difference test (pLDDT). High-confidence regions are shown in blue, while lower-confidence regions are indicated by shades of yellow to orange. **b**, The SDS-PAGE is shown the final protein purify of HTLV-1 CA_full-length_, CA_NTD_ and CA_CTD_.

**Extended Data Fig. 3.**
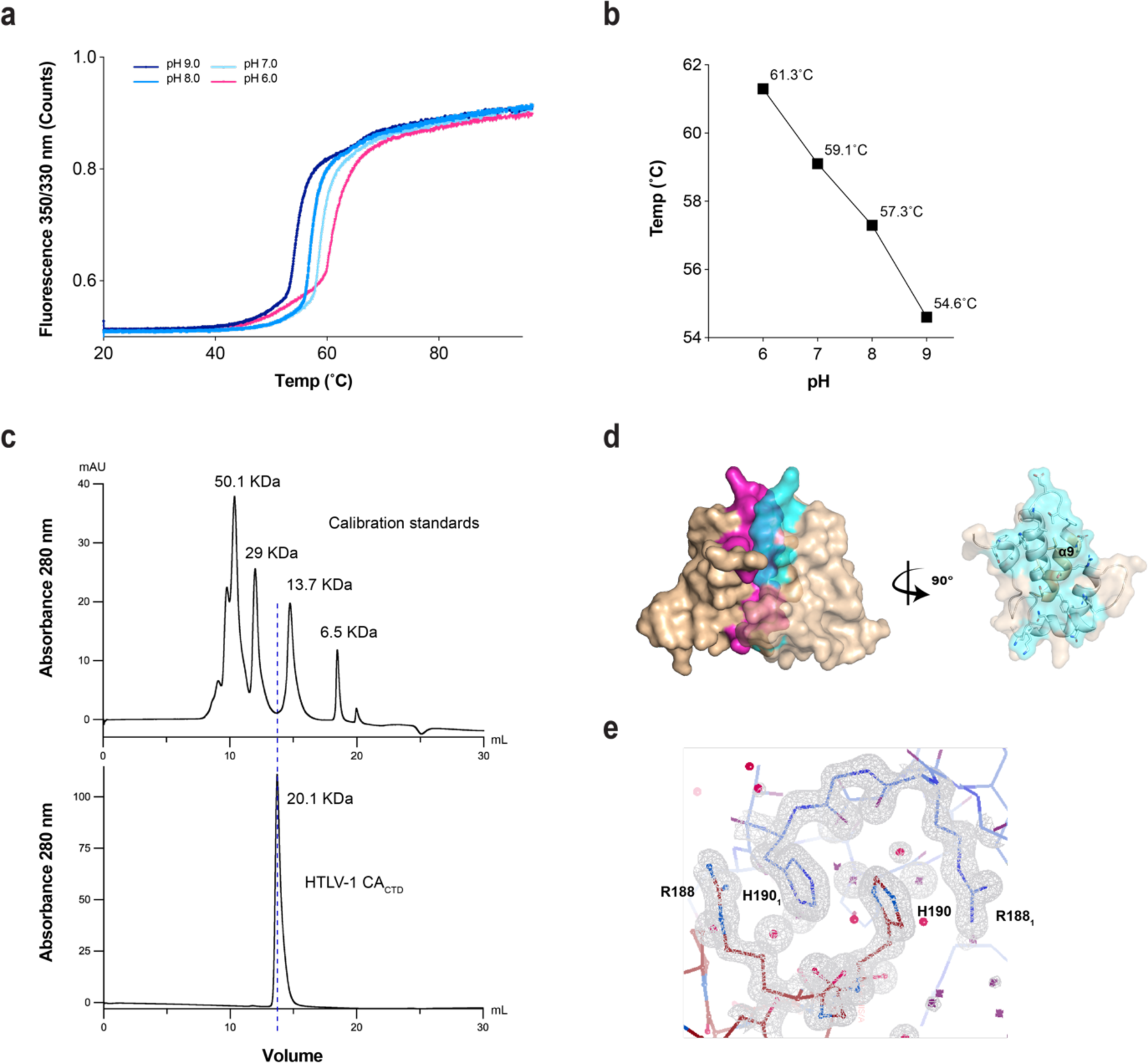
HTLV-1 CA thermal stability and CA_CTD_ dimer characters. **a** and **b**, Graphs demonstrate the pH-dependent thermal unfolding traces of HTLV-1 CA_full-length_ protein (**a**), highlighting unfolding temperature points corresponding to pH 6 to 9 (**b**). **c**, The aligned elution curves from size-exclusion chromatography display the calibration standards (top) and a dimer size of HTLV-1 CA_CTD_ (down). **d**, The CA_CTD_ dimer interface (magenta and cyan) is driven by α-helix 9 showed in surface. **e**, The CA_CTD_ residues R188 and H190 (in red) interdigitate with R188_1_ and H190_1_ (in blue) from another copy of molecule generated by crystallographic symmetry.

**Extended Data Fig. 4.**
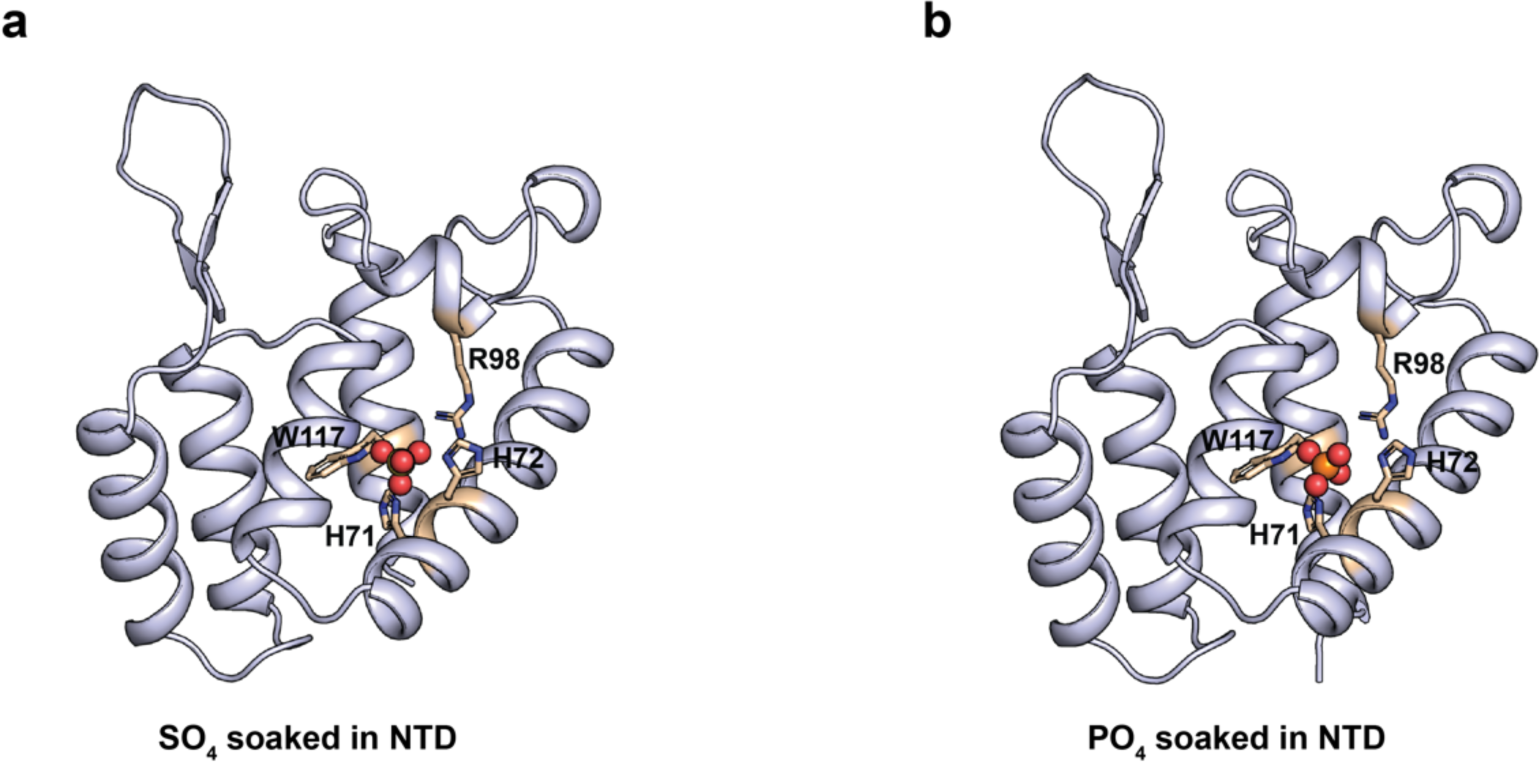
The solved sulfate-bound and phosphate-bound structures within the CA_NTD_ triclinic crystal form. A sulfate ion (**a**) or a phosphate ion (**b**) was soaked in the cross-linked CA_NTD_ triclinic crystal form, coordinating with residues H71, H72, R98 and W117.

**Extended Data Fig. 5.**
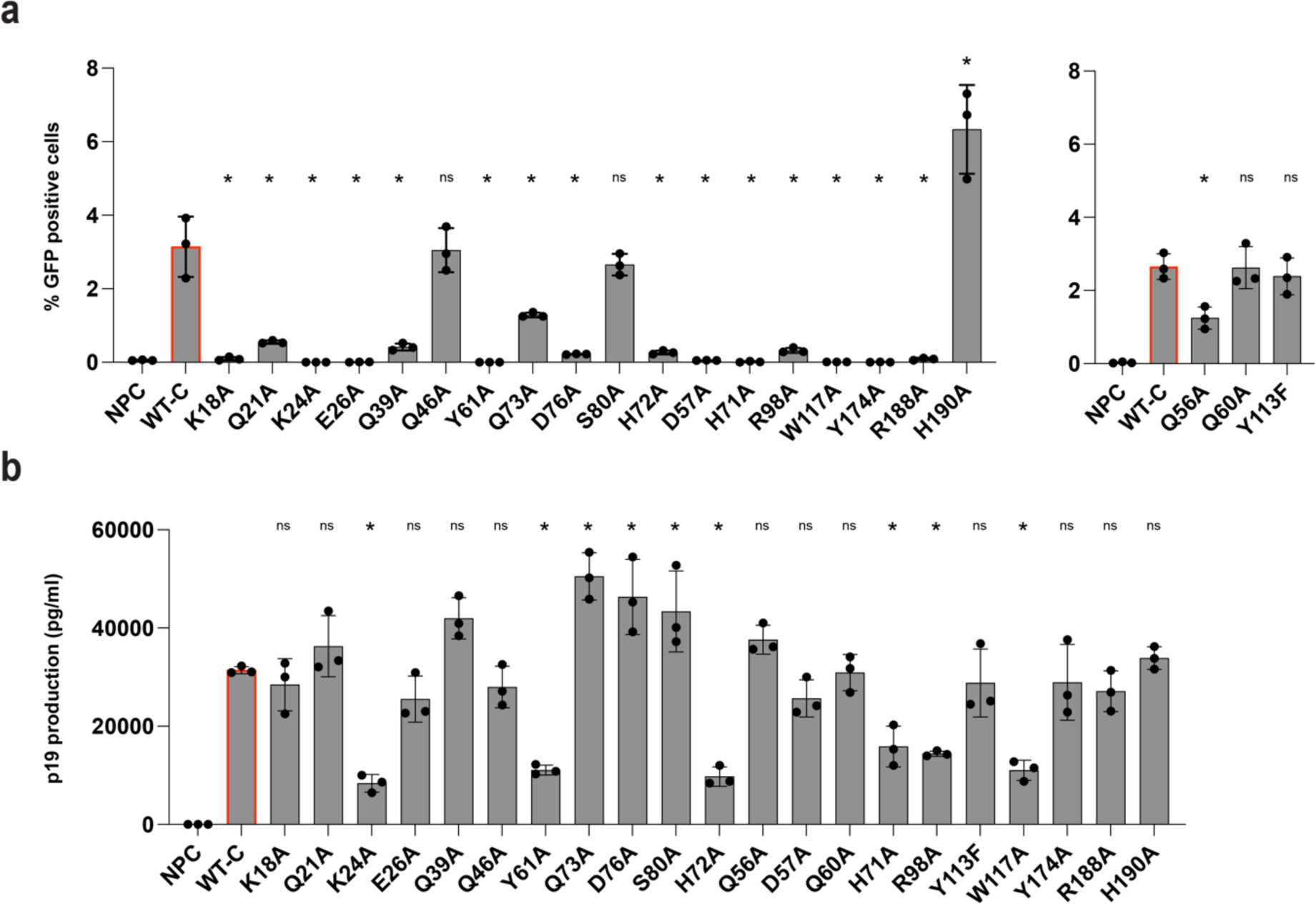
Infectivity assays of HTLV-1 mutants. **a**, Bar graph displaying infectivity of WT-C CA mutants as measured by GFP reporter expression in a replication dependent infection system. **b**, p19 production as measured using a p19 ELISA. Results are displayed as the average of 3 independent experiments with error bars representing the SD. Statistical analysis was performed using a one-way ANOVA with comparison to the wild-type control (WT-C, highlighted with a red border). * = p < 0.05, *ns* = p > 0.05.

**Table S1.**
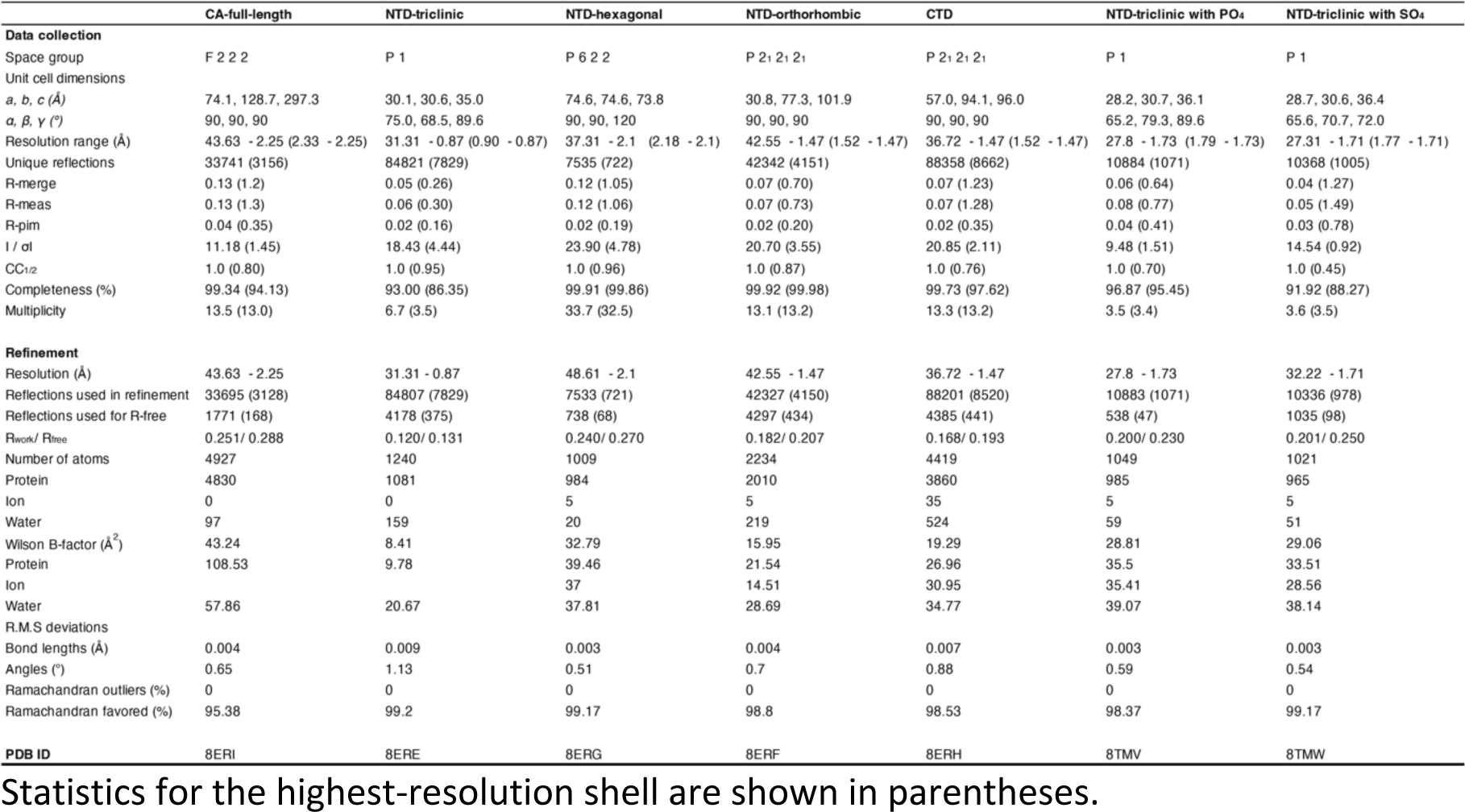
Data collection and refinement statistics.

## Materials and methods

### HTLV-1 CA constructs design and cloning

HTLV-1, Aus-GM variant of C type, CA full-length protein consists of 214 amino acids with 23.8 kDa in molecular weight. Previous work reported that the CA protein precipitated during purification (15). To enhance the homogeneous of CA, we designed constructs based on an AlphaFold2 predicted model, showing a flexible hinge as the domain boundary between NTD and CTD. Consequently, three constructs were generated: (1) full-length protein, HTLV-1 CA, comprising resides of P1-L214; (2) N-terminal domain, HTLV-1 CA_NTD_, comprising residues of P1-S127 that includes the intact N-terminal β-hairpin; and (3) C-terminal domain, HTLV-1 CA_CTD_, comprising residues of A128-L214. The gene was synthesised and subcloned into the pET-30a (+) respectively by Genscript.

### Protein overexpression and purification

Plasmids of HTLV-1 CA, CA_NTD_ and CA_CTD_ were transformed in *E.coli* Rosetta 2 (DE3) pLysS cells. A single colony was picked and grown in 10 mL LB medium with 30 μg/L Kanamycin and 34 μg/L Chloramphenicol at 37 °C. The starter culture was then used to inoculate 1 L of LB medium at a 1:100 ratio, inducing cells with 1 mM IPTG when OD_600_ reached to 0.6 and incubating at 30°C for 4 h (180 rpm). Cells were harvested by centrifugation (6000 g, 15 min, 4 °C) and lysed by sonication in precooled lysis buffer, containing 50 mM Tris-HCl, pH 7.0, 50 mM NaCl, 20 mM β-mercaptoethanol and a tablet of the complete EDTA-free Protease Inhibitor (Roche). The cell debris was removed by centrifugation (29,300 g, 30 min, 4°C) and the soluble portion was collected for further purification.

HTLV-1 CA was further purified by ammonium sulfate (20% w/v) precipitation, with solution stirred 30 min at 4°C and spined down (29,300 g, 20 min, 4°C). The pellet was resuspended in 100 mM citric acid (pH 5.0) with 20 mM β-mercaptoethanol, and dialysed against 2 L of the same buffer for at least three buffer changes. Before ion-exchange chromatography, the dialysed protein was briefly centrifuged (29,300 g, 20 min, 4°C) and the soluble fraction was passed through a 0.45 µM syringe filter (Millipore, Merk) before loading onto HiLoad 16/10 SP Sepharose HP column (Cytiva), running with buffer A, 50 mM citric acid (pH 5.0) and buffer B, 50 mM citric acid (pH 5.0) supplied with 1 M NaCl. The expected eluent was dialysed against 20 mM HEPES (pH 7.0) with 40 mM NaCl and briefly centrifuged (29,300 g, 20 min, 4°C). The soluble portion was passed through a 0.45 µM syringe filter (Millipore, Merk) and loaded onto a HiLoad 16/60 Superdex 200 pg (Cytiva) column for size-exclusion chromatography (SEC) running with the same buffer condition as for dialysis. The expected fractions were concentrated to 12 mg/mL, aliquoted and frozen in liquid nitrogen for storage at −80°C.

HTLV-1 CA_NTD_ protein was purified by following the same protocol of ammonium sulfate precipitation and ion-exchange chromatography for CA full-length. It was purified onto HiLoad 16/60 Superdex 75 pg (Cytiva) SEC column. The purified fractions were concentrated to 16 mg/mL, aliquoted and frozen in liquid nitrogen for storage at −80°C.

HTLV-1 CA_CTD_ protein was purified further through two-step ammonium sulfate precipitation. Firstly, the junk aggregates were precipitated by 20% w/v ammonium sulfate and removed by centrifugation (29,300 g, 20 min, 4°C). The target protein retained in the soluble fraction was reprecipitated with 30% w/v ammonium sulfate, pelleted by centrifugation (29,300 g, 20 min, 4°C), and resuspended in 100 mM citric acid (pH 5.0) with 20 mM β-mercaptoethanol for dialysis with at least three buffer changes. The remaining steps followed the purification protocol of HTLV-1 CA_NTD_. The purified protein was concentrated to 40 mg/mL, aliquoted and frozen in liquid nitrogen for storage at −80°C.

### Protein crystallisation and X-ray diffraction

All the protein crystals were initially screened by sitting-drop vapour diffusion method using a crystallisation robot (NT8, Formulatrix) combined with a dispenser for high-throughput preparation of crystal trays (Art Robbins Crystal Phoenix). 100 nL protein was mixed with 100 nL precipitant (commercial kits) on the 96-well tray (MiTeGen), evaporating against 40 μL reservoir precipitant at 20°C. The screened crystals were manually refined by either sitting- or hanging-drop vapour diffusion method.

The crystal of HTLV-1 CA was optimised by seeding microcrystals into a 4 μL hanging-drop mixture, composed of 2 μL protein solution (5 mg/mL) and 2 μL precipitant (8% v/v 2-propanol, 23% v/w PEG 3350, 0.1 M sodium citrate, pH 6.0), equilibrating against 500 μL of reservoir solution at 20°C. The optimised crystals were cryoprotected in the mother liquor.

HTLV-1 CA_NTD_ protein was crystallised in three crystal forms that were optimised by sitting-drop vapour diffusion at 20°C. A 2 µL drop was prepared with 1 μL protein (10 mg/mL) and 1 μL precipitant, and equilibrating against 80 μL reservoir solution. The precipitants used for the three crystal forms were: 1) triclinic, 10% ethylene glycol, 10% PEG 8K, 0.1 M HEPES, pH 7.5; 2) orthorhombic, 0.2 M (NH_4_)_2_SO_4_, 25% PEG 4000, 0.1 M Na acetate, pH 4.6; and 3) hexagonal, 2 M (NH_4_)_2_SO_4_, 2% PEG 400, 0.1 M HEPES, pH 7.5. The cryoprotectants for each crystal form were prepared by supplementing its respective mother liquor with 20% ethylene glycol, 20% MPD, and 10% glycerol, respectively.

The crystals of HTLV-1 CA_CTD_ were also optimised by sitting-drop vapour diffusion method at 20°C. A 2 µL drop was prepared with 1 μL protein (20 mg/mL) and 1 μL precipitant (1.5 M (NH_4_)_2_SO_4_, 0.1 M Na_2_HPO_4_, pH 4.2 adjusted with citric acid), and equilibrating against 80 μL of reservoir solution. Crystals were cryoprotected in mineral oil.

All cryoprotected crystals were flash-frozen in liquid nitrogen for X-ray diffraction.

### Triclinic crystal cross-linking and soaking

The crystal cross-linking reactions were performed in a 24-well tray with each well containing 0.5 mL crystallising solution and a micro-bridge serving as an isolated reservoir for 2 μL of 25% glutaraldehyde (pH 3.0). A cover slip, hanging a droplet with crystals, was sealed over the top of the well for overnight. The cross-linked crystals were briefly washed with mother liquor to stop cross-link reaction, then soaked into 2 M ammonium sulfate (pH 6.0) or 2 M sodium/potassium phosphate (pH 6.0) for 4 h. Lastly, crystals were cryoprotected in the mother solution with 20% glycerol before flash freezing in liquid nitrogen for X-ray diffraction.

### Data collection, structure solution and refinement

All datasets were remotely collected and auto processed at MX2 beamline (wavelength 0.9537, energy 13 keV) of the Australian Synchrotron^47^. For collecting ultra-high resolution datasets of HTLV-1 CA_NTD_ triclinic crystals, the wavelength was adjusted to 0.7999 Å for a higher energy of 15.5 keV. The structures of HTLV-1 CA_NTD/CTD_ were determined by utilising AlphaFold2 predicted models as templates for molecular replacement in Phenix (Version 1.20.1-4487) that was also applied for subsequent rounds of refinement along with Coot. The HTLV-1 CA full-length structure was solved by molecular replacement using the solved CA_NTD_ and CA_CTD_ as searching models. The quality of determined structures was accessed for Ramachandran outliers, incorrect stereochemistry, rotamer outliers and steric clashes through MOLPROBITY. All structural figures were rendered in PyMOL Molecular Graphics System (Version 2.5.3 Schrödinger). The final statistics of all data collections and refinements are presented in Table S1.

### High resolution protein thermal stability analysis

Measurements were performed using a nanoDSF (differential scanning fluorimetry) instrument (Prometheus NT.48), which detects subtle changes in the fluorescence (330/350 nm) of the tryptophan of proteins, offering a sensitive label-free monitoring for protein unfolding and refolding processes. This approach allows a rapid screening for optimal protein purification and storage conditions.

The samples were prepared at a final concentration of 3 mg/ml HTLV-1 CA protein in pH 6 (50 mM MES), 7 (50 mM HEPES), 8 (50 mM TRIS) and 9 (50 mM TRIS), respectively, up to a total volume of 20 µL. By capillarity action, each sample was aspirated into standard-treated glass capillaries (NanoTemper) that were loaded into a 48-capillary sample holder. The excitation wavelength was set at 280 nm (20 nm bandwidth), and the LED power was adjusted to produce emission intensities between 330 and 380 nm, ranging from 6000 to 25,000 A.U. The intrinsic fluorescence was measured for 1 to 5 s in the absence of a heat gradient, setting the temperature varying from 20 to 90°C.

### HTLV-1 CA mutational analysis – infection and viral phenotype

The C-type CA sequence was cloned into pCMVHTLV_delta env (kind gift from Dimitry Mazurov) by Gibson Assembly (NEB) and all subsequent point mutations were introduced by standard overlapping PCR. The sequences of the primers used to generate the mutations are provided in Supplementary methods. The DNA sequences of all constructs were verified by whole plasmid sequencing (Primordium Inc).

HEK-293T (ATCC) cells were maintained in Dulbecco’s Modified Eagle Medium (DMEM) with high glutamine and glucose, supplemented with 10% FCS. Cells were passaged every 2 days and maintained at 37°C with 5% CO_2_. Cells were routinely tested for mycoplasma contamination.

HEK-293T cells were plated at 2.5×10^5^ cells/well in 12 well plates the day before transfection. Transfections were performed using pCMVHTLV-1_delta env, pCRU5-inGFPt (Addgene #60236)) and pVSV-G at a 1:1:0.5 ratio using Fugene HD. For HIV infections, the following plasmids were used for transfections; psPAX2, pUCHR-inGFPt (Addgene #60237) and pVSV-G. Media was replaced 16 hours post transfection. For lenacapavir (MedChemExpress) experiments, indicated concentrations were added 30 mins prior to transfection and again during media replacement. 72 hours post transfection, cells and supernatants were harvested for analysis of infectivity and virus production, respectively. For infectivity measurements, cells were stained with Live/Dead Fixable Near-IR Dead Cell Stain (ThermoFisher) for 20 mins at RT and then fixed in 2% paraformaldehyde (Electron Microscopy Sciences) for 20 mins at RT. Samples were run on a BD LSR Fortessa X-20 and the number of GFP positive cells enumerated using FlowJo V10.

Virus production was measured using a p19 antigen ELISA kit (Zeptometrix) according to manufacturer’s instructions.

## Acknowledgements

We thank C. Dickson for critical reading of this manuscript. This work was supported by a National Health and Medical Research Council Ideas Grant (GNT2013215; D.A.J., T.B.) and Wellcome Trust Collaborator Award (214344/Z/18/Z; D.A.J., T.B.). D.A.J. was supported by a UNSW Scientia Fellowship. We also acknowledge the Structural Biology Facility in the Mark Wainwright Analytical Centre – UNSW, funded in part by the Australian Research Council Linkage Infrastructure, Equipment and Facilities Grant: ARC LIEF 190100165. This research was undertaken in part using the MX2 beamline at the Australian Synchrotron, part of ANSTO, and made use of the Australian Cancer Research Foundation (ACRF) detector. We acknowledge the Bedegal people of the Eora nation, the traditional custodians of the land upon which this research took place.

## Author contributions

R.Y. was responsible for construct design, recombinant protein production, crystallization, X-ray data collection, and structure solution with the support of N.L. P.P. adapted the single-cycle infection model for the study of subtype C capsid, and performed all infection-relevant experiments. T.B. and D.A.J. conceptualized the project. R.Y. and D.A.J. wrote the manuscript with input from all authors.

## Author information

The authors declare no competing financial interests. Correspondence and requests for materials should be addressed to D.A.J.

## Reference

1. Tagaya, Y. & Gallo, R.C. The Exceptional Oncogenicity of HTLV-1. Front Microbiol 8, 1425 (2017).

2. Einsiedel, L., Cassar, O., Bardy, P., Kearney, D. & Gessain, A. Variant human T-cell lymphotropic virus type 1c and adult T-cell leukemia, Australia. Emerg Infect Dis 19, 1639–41 (2013).

3. Kaplan, J.E. et al. The risk of development of HTLV-I-associated myelopathy/tropical spastic paraparesis among persons infected with HTLV-I. J Acquir Immune Defic Syndr (1988) 3, 1096–101 (1990).

4. Bangham, C.R., Araujo, A., Yamano, Y. & Taylor, G.P. HTLV-1-associated myelopathy/tropical spastic paraparesis. Nat Rev Dis Primers 1, 15012 (2015).

5. Einsiedel, L. et al. Clinical associations of Human T-Lymphotropic Virus type 1 infection in an indigenous Australian population. PLoS Negl Trop Dis 8, e2643 (2014).

6. Einsiedel, L., Fernandes, L., Spelman, T., Steinfort, D. & Gotuzzo, E. Bronchiectasis is associated with human T-lymphotropic virus 1 infection in an Indigenous Australian population. Clin Infect Dis 54, 43–50 (2012).

7. Martin, F., Taylor, G.P. & Jacobson, S. Inflammatory manifestations of HTLV-1 and their therapeutic options. Expert Rev Clin Immunol 10, 1531–46 (2014).

8. LaGrenade, L., Hanchard, B., Fletcher, V., Cranston, B. & Blattner, W. Infective dermatitis of Jamaican children: a marker for HTLV-I infection. Lancet 336, 1345–7 (1990).

9. Tagaya, Y., Matsuoka, M. & Gallo, R. 40 years of the human T-cell leukemia virus: past, present, and future. F1000Res 8(2019).

10. Afonso, P.V., Cassar, O. & Gessain, A. Molecular epidemiology, genetic variability and evolution of HTLV-1 with special emphasis on African genotypes. Retrovirology 16, 39 (2019).

11. Gessain, A. & Cassar, O. Epidemiological Aspects and World Distribution of HTLV-1 Infection. Front Microbiol 3, 388 (2012).

12. Einsiedel, L. et al. Very high prevalence of infection with the human T cell leukaemia virus type 1c in remote Australian Aboriginal communities: Results of a large cross-sectional community survey. PLoS Negl Trop Dis 15, e0009915 (2021).

13. Cassar, O., Einsiedel, L., Afonso, P.V. & Gessain, A. Human T-cell lymphotropic virus type 1 subtype C molecular variants among indigenous australians: new insights into the molecular epidemiology of HTLV-1 in Australo-Melanesia. PLoS Negl Trop Dis 7, e2418 (2013).

14. Pontillo, A., Girardelli, M., Catamo, E., Duarte, A.J. & Crovella, S. Polymorphisms in TREX1 and susceptibility to HIV-1 infection. Int J Immunogenet 40, 492–4 (2013).

15. Zuliani-Alvarez, L. et al. Evasion of cGAS and TRIM5 defines pandemic HIV. Nat Microbiol 7, 1762–1776 (2022).

16. Stephens, C. & Naghavi, M.H. The host cytoskeleton: a key regulator of early HIV-1 infection. FEBS J (2022).

17. Jacques, D.A. et al. HIV-1 uses dynamic capsid pores to import nucleotides and fuel encapsidated DNA synthesis. Nature 536, 349–53 (2016).

18. Dickson, C.F. et al. The HIV capsid mimics karyopherin engagement of FG-nucleoporins. Nature 626, 836–842 (2024).

19. Fu, L. et al. HIV-1 capsids enter the FG phase of nuclear pores like a transport receptor. Nature 626, 843–851 (2024).

20. Zila, V. et al. Cone-shaped HIV-1 capsids are transported through intact nuclear pores. Cell 184, 1032–1046.e18 (2021).

21. Scoca, V., Morin, R., Collard, M., Tinevez, J.Y. & Di Nunzio, F. HIV-induced membraneless organelles orchestrate post-nuclear entry steps. J Mol Cell Biol 14(2023).

22. Ganser, B.K., Li, S., Klishko, V.Y., Finch, J.T. & Sundquist, W.I. Assembly and analysis of conical models for the HIV-1 core. Science 283, 80–3 (1999).

23. Zhang, M.J., Stear, J.H., Jacques, D.A. & Böcking, T. Insights into HIV uncoating from single-particle imaging techniques. Biophys Rev 14, 23–32 (2022).

24. Rihn, S.J. et al. Extreme genetic fragility of the HIV-1 capsid. PLoS Pathog 9, e1003461 (2013).

25. Paik, J. Lenacapavir: First Approval. Drugs 82, 1499–1504 (2022).

26. Mushtaq, A. & Kazi, F. Lenacapavir: a new treatment of resistant HIV-1 infections. Lancet Infect Dis 23, 286 (2023).

27. Bekker, L.G. et al. Twice-Yearly Lenacapavir or Daily F/TAF for HIV Prevention in Cisgender Women. N Engl J Med (2024).

28. Link, J.O. et al. Clinical targeting of HIV capsid protein with a long-acting small molecule. Nature 584, 614–618 (2020).

29. Rebensburg, S.V. et al. Sec24C is an HIV-1 host dependency factor crucial for virus replication. Nat Microbiol 6, 435–444 (2021).

30. Price, A.J. et al. Host cofactors and pharmacologic ligands share an essential interface in HIV-1 capsid that is lost upon disassembly. PLoS Pathog 10, e1004459 (2014).

31. Faysal, K.M.R. et al. Pharmacologic hyperstabilisation of the HIV-1 capsid lattice induces capsid failure. Elife 13(2024).

32. Schaller, T. et al. HIV-1 capsid-cyclophilin interactions determine nuclear import pathway, integration targeting and replication efficiency. PLoS Pathog 7, e1002439 (2011).

33. Tang, C., Ndassa, Y. & Summers, M.F. Structure of the N-terminal 283-residue fragment of the immature HIV-1 Gag polyprotein. Nat Struct Biol 9, 537–43 (2002).

34. Gres, A.T., et al. STRUCTURAL VIROLOGY. X-ray crystal structures of native HIV-1 capsid protein reveal conformational variability. Science 349, 99–103 (2015).

35. Mattei, S., Glass, B., Hagen, W.J., Kräusslich, H.G. & Briggs, J.A. The structure and flexibility of conical HIV-1 capsids determined within intact virions. Science 354, 1434–1437 (2016).

36. Mallery, D.L. et al. IP6 is an HIV pocket factor that prevents capsid collapse and promotes DNA synthesis. Elife 7(2018).

37. Khorasanizadeh, S., Campos-Olivas, R. & Summers, M.F. Solution structure of the capsid protein from the human T-cell leukemia virus type-I. J Mol Biol 291, 491–505 (1999).

38. Zhao, G. et al. Mature HIV-1 capsid structure by cryo-electron microscopy and all-atom molecular dynamics. Nature 497, 643–6 (2013).

39. Stacey, J.C.V. et al. Two structural switches in HIV-1 capsid regulate capsid curvature and host factor binding. Proc Natl Acad Sci U S A 120, e2220557120 (2023).

40. Shunaeva, A. et al. Improvement of HIV-1 and Human T Cell Lymphotropic Virus Type 1 Replication-Dependent Vectors via Optimization of Reporter Gene Reconstitution and Modification with Intronic Short Hairpin RNA. J Virol 89, 10591–601 (2015).

41. Kalemera, M.D., Maher, A.K., Dominguez-Villar, M. & Maertens, G.N. Cell Culture Evaluation Hints Widely Available HIV Drugs Are Primed for Success if Repurposed for HTLV-1 Prevention. Pharmaceuticals (Basel) 17(2024).

42. Kelly, B.N. et al. Structure of the antiviral assembly inhibitor CAP-1 complex with the HIV-1 CA protein. J Mol Biol 373, 355–66 (2007).

43. Gamble, T.R. et al. Structure of the carboxyl-terminal dimerization domain of the HIV-1 capsid protein. Science 278, 849–53 (1997).

44. Obr, M. et al. Distinct stabilization of the human T cell leukemia virus type 1 immature Gag lattice. Nat Struct Mol Biol (2024).

45. Smyth, M., Pettitt, T., Symonds, A. & Martin, J. Identification of the pocket factors in a picornavirus. Arch Virol 148, 1225–33 (2003).

46. Robert, X. & Gouet, P. Deciphering key features in protein structures with the new ENDscript server. Nucleic Acids Res 42, W320–4 (2014).

47. McPhillips, T.M. et al. Blu-Ice and the Distributed Control System: software for data acquisition and instrument control at macromolecular crystallography beamlines. J Synchrotron Radiat 9, 401–6 (2002).

